# How Do Deer Respiratory Epithelial Cells Weather The Initial Storm of SARS-CoV-2?

**DOI:** 10.1101/2023.04.24.538130

**Authors:** Kaitlyn M. Sarlo Davila, Rahul K. Nelli, Kruttika S. Phadke, Rachel M. Ruden, Sang Yongming, Bryan H. Bellaire, Luis G. Gimenez-Lirola, Laura C. Miller

## Abstract

The potential infectivity of SARS-CoV-2 in animals raises a public health and economic concern, particularly the high susceptibility of white-tailed deer (WTD) to SARS-CoV-2. The disparity in the disease outcome between humans and WTD is very intriguing, as the latter are often asymptomatic, subclinical carriers of SARS-CoV-2. To date, no studies have evaluated the innate immune factors responsible for the contrasting SARS-CoV-2-associated disease outcomes in these mammalian species. A comparative transcriptomic analysis in primary respiratory epithelial cells of human (HRECs) and WTD (Deer-RECs) infected with SARS-CoV-2 was assessed throughout 48 hours post inoculation (hpi). Both HRECs and Deer-RECs were susceptible to SARS-COV-2, with significantly (*P* < 0.001) lower virus replication in Deer-RECs. The number of differentially expressed genes (DEG) gradually increased in Deer-RECs but decreased in HRECs throughout the infection. The ingenuity pathway analysis of DEGs further identified that genes commonly altered during SARS-CoV-2 infection mainly belong to cytokine and chemokine response pathways mediated via IL-17 and NF-κB signaling pathways. Inhibition of the NF-κB signaling in the Deer-RECs pathway was predicted as early as 6 hpi. The findings from this study could explain the lack of clinical signs reported in WTD in response to SARS-CoV-2 infection as opposed to the severe clinical outcomes reported in humans.

**HIGHLIGHTS:**

1. White-tailed deer primary respiratory epithelial cells are susceptible to SARS- CoV-2 without causing hyper cytokine gene expression.
2. Downregulation of IL-17 and NF-κB signaling pathways after SARS-CoV-2 infection could be key to the regulated cytokine response in deer cells.
3. Deer innate immune system could play a critical role in early antiviral and tissue repair response following SARS-CoV-2 infection.

## MAIN

Deer hunting and sales of captive deer contribute >$20 billion to the US GDP directly and indirectly, and support over >300k jobs associated with these industries ^1, 2^. The human interaction with deer in the US is relatively high, with nearly 8 million people spending over 115 million days in the field for deer hunting in 2016 ^1^. During the same year, an astonishing 30.1 million individuals, almost one-tenth of the U.S. population, engaged in watching wild mammals like deer near their homes, meanwhile another 14.5 million individuals reported feeding non-avian wildlife ^3^. Importantly, this overlooks the possibilities of more intimate and sustained human-deer interaction through wildlife rehabilitation or captive settings and fails to account for the full extent of time people spent in natural habitats engaged in other forms of outdoor recreation. This widespread human-deer interaction creates a significant risk of exposure to the North American white-tailed Deer (*Odocoileus virginianus*; WTD) for diseases like SARS-CoV-2, which causes COVID-19.

The high susceptibility of WTD to SARS-CoV-2 infection, their ability to transmit the virus to other Deer ^4–10^, and the potential for spillback to humans can have significant health and economic consequences. Further studies are warranted to better understand the infection and transmission dynamics of SARS-CoV-2 in WTD, and these studies would be crucial in developing appropriate mitigation strategies and minimizing the risk of spillback to humans. Cumulative evidence suggests that subclinical infection and asymptomatic carriage of SARS-CoV-2 are common in WTD ^7, 8, 10, 11^. Experimental infection studies in WTD have shown SARS-CoV-2 infection rates of up to 40%, along with shedding and transmitting the virus for up to five days post-infection ^5, 6, 9^. High levels of viremia and virus shedding have been reported in deer, which could lead to environmental or aerosol transmission ^8–12^. However, no reports of a clinical illness associated with SARS-CoV-2 in the deer populations surveyed, and experimental conditions studies reported only subclinical infections in white-tailed deer challenged with SARS-CoV-2 ^8, 11, 13^. Contrastingly, in most human cases, SARS-CoV-2 cause subclinical to mild disease, but a significant number of cases develop severe symptoms that can lead to long-lasting lung damage or death ^14–16^. These severe cases are often associated with high levels of proinflammatory cytokines and low antiviral responses, leading to systemic complications ^15, 17, 18^.

SARS-CoV-2 replicates in the upper respiratory tract of both humans and deer ^8, 10, 19^, which would justify using primary respiratory epithelial cell cultures derived from WTD as *in vitro* infection model to evaluate cell-virus interactions during SARS-CoV-2 infection on a daily/hourly basis and under controlled conditions. In addition, studies have demonstrated the susceptibility of human respiratory epithelial cells (HRECs) to SARS-CoV-2 infection ^20, 21^. In the current study, SARS-CoV-2 infection studies were performed using primary WTD respiratory epithelial cells (Deer-RECs) and HRECs. To determine early cell-virus interactions in these cell types derived from the hosts with contrasting disease outcomes, a comparative transcriptome-wide analysis was performed using RNA-Seq analysis.

## RESULTS

### Both Deer-RECs and HRECs are susceptible to SARS-CoV-2 infection

Deer-RECs and HRECs cultures were inoculated with six different viral doses (10^5^, 10^4^, 10^3^, 10^2^, 10, 1 PFU/mL) and corresponding mock-inoculated controls. Cells were incubated and monitored daily for 120 hpi. Virus-specific CPE, such as rounding of cells, vacuolation, and cell detachment/death, were observed at 48 hpi in Deer-RECs at doses >10^3^ PFU/mL, while in HRECs, CPE was evident by 72 hpi. Mock-inoculated controls showed no CPE. The CPE was time and virus-dose-dependent in both Deer- RECs and HRECs. However, cell detachment/cell death was remarkably higher in Deer- RECs compared to HRECs at viral doses ≥ 10^2^ PFU/mL between 48-120 hpi.

Microscopy findings were further supported by ICC staining for SARS-CoV-2 N protein (stained brown), indicating viral replication and active production of viral proteins in HRECs (Fig. 1A) and Deer-RECs (Fig. 1B) inoculated with SARS-CoV-2.

**Figure 1.**
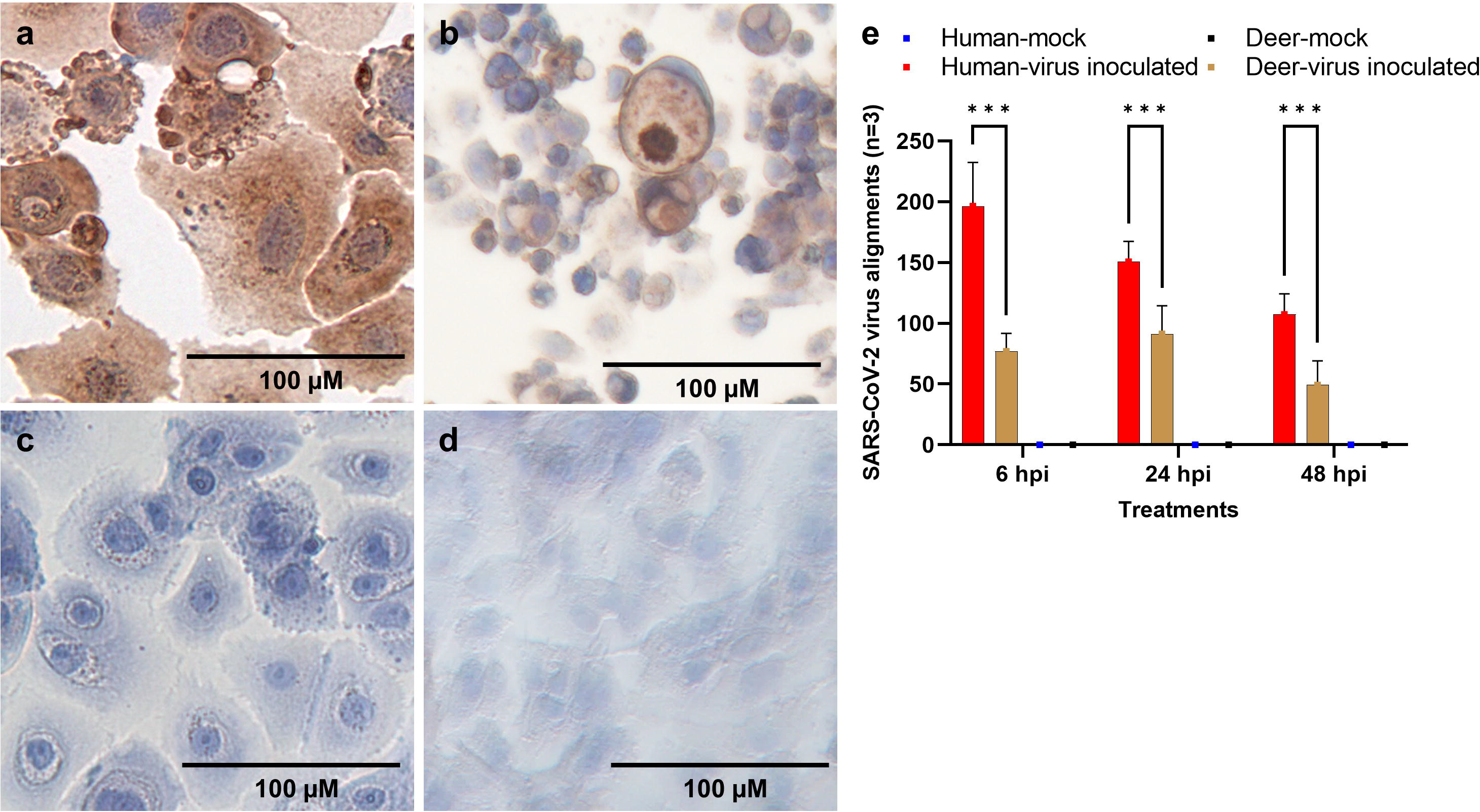
Detection of SARS-CoV-2 infection in primary respiratory epithelial cells of human and white-tailed deer inoculated with SARS-CoV-2 virus isolate USA- WA1/2020. (a-d) Cells fixed in 4% paraformaldehyde were stained for SARS-CoV-2 viral N protein with ImmPRESS VR anti-rabbit IgG horseradish peroxidase (HRP) polymer detection kit (MP-6401-15; Vector Laboratories) and a recombinant anti-SARS- CoV-2 N protein rabbit monoclonal antibody (0.75 μg/mL) [The following reagent was obtained through BEI Resources, NIAID, NIH: Monoclonal Anti-SARS Coronavirus/SARS-Related Coronavirus 2 Nucleocapsid Protein (produced in vitro), NR-53791; SinoBio Cat: 40143-R001]. Dark brown represents a positive antibody expression, pale brown represents background staining, and blue represents the nucleus counterstained with hematoxylin; n=6 and scale bar-100_μm. Cells inoculated with a viral dose of 10^5^ PFU/mL (a-Human; b- Deer) or culture medium (c- Human; d- Deer). (e) Bar graph showing the average number of viral alignment counts of SARS- CoV-2 reference genomes across different time points in human (red) and deer (brown) in respiratory epithelial cells. Data represent three technical replicates with SEM as error bars. *** denotes *P* < 0.001

Correspondingly, mock-inoculated HRECs (Fig. 1C) and Deer-RECs (Fig. 1D) remained negative throughout the observation period. Interestingly, the cellular nucleus remained intact in Deer-RECs and HRECs stained with hematoxylin in both infected and mock-inoculated control cells.

Based on the dose-response data of virus-induced CPE and ICC, a viral dose of 10^2^ PFU/mL was selected for gene expression analysis of the early innate immune response in Deer-RECs and HRECs at 6, 24, and 48 hpi. In addition to the CPE results, the susceptibility of Deer-RECs and HRECs cultures to virus infection was further confirmed by transcriptomic alignments to the SARS-CoV-2 reference genome. No sequence from mock-inoculated culture samples aligned to the virus genome, but several alignments were found in the virus-inoculated samples. The average number of viral sequence alignments for the three samples for each time-point and species were shown in Fig. 1E.

### Differential gene expression in Deer-RECs and HRECs infected with SARS-CoV-2

The total RNA from Deer-RECs and HREC virus- and mock-inoculated culture samples sequenced had RNA Integrity Numbers (RIN) ranging from 9.7 to 10, and ∼5,000,000 reads per sample were generated from the sequencing. Volcano plots generated using DEG data from SARS-CoV-2 infected vs. corresponding mock controls in HREC and Deer-REC samples show upregulated genes in red and downregulated genes in green for each time point (Fig. 2a). In HRECs, there was a gradual decrease in the number of DEGs with the progression of infection, and a high number of DEGs were observed as early as 6 hpi (491 DEGs; 394 upregulated; 97 downregulated) followed by 24 hpi (123 DEGs; 23 upregulated;100 downregulated), and 48 hpi (70 DEGs; 36 upregulated; 34 downregulated). In contrast, the number of significant DEGs increased over the course of the infection in Deer-RECs, where 36 DEGs were significant at 6 hpi (29 upregulated; 7 downregulated), 135 at 24 hpi (99 upregulated; 34 downregulated), and 280 at 48 hpi (134 upregulated; 146 downregulated); for additional information, refer to Supplementary Data. To delineate the shared or uniquely expressed DEGs, the data was analyzed using multidimensional six-set Venn diagrams showing upregulated and downregulated DEGs (shown separately) shared between species and time points (Fig. 2b). At 6 hpi, only eight genes were commonly upregulated between both species, while 372 genes were unique in HRECs. At 24 hpi, three in common and 18 genes unique to HRECs, while at 48 hpi, only one in common but 33 genes unique to HRECs and 113 genes to Deer-RECs. In the case of downregulated genes, both species did not share any genes at 6 hpi, while two and one genes were shared at 24 and 48 hpi, respectively. In Deer-RECs, 25, 19, and 113 genes were downregulated, while in HRECs, 85, 71, and 24 were downregulated at 6, 24, and 48 hpi, respectively.

**Figure 2.**
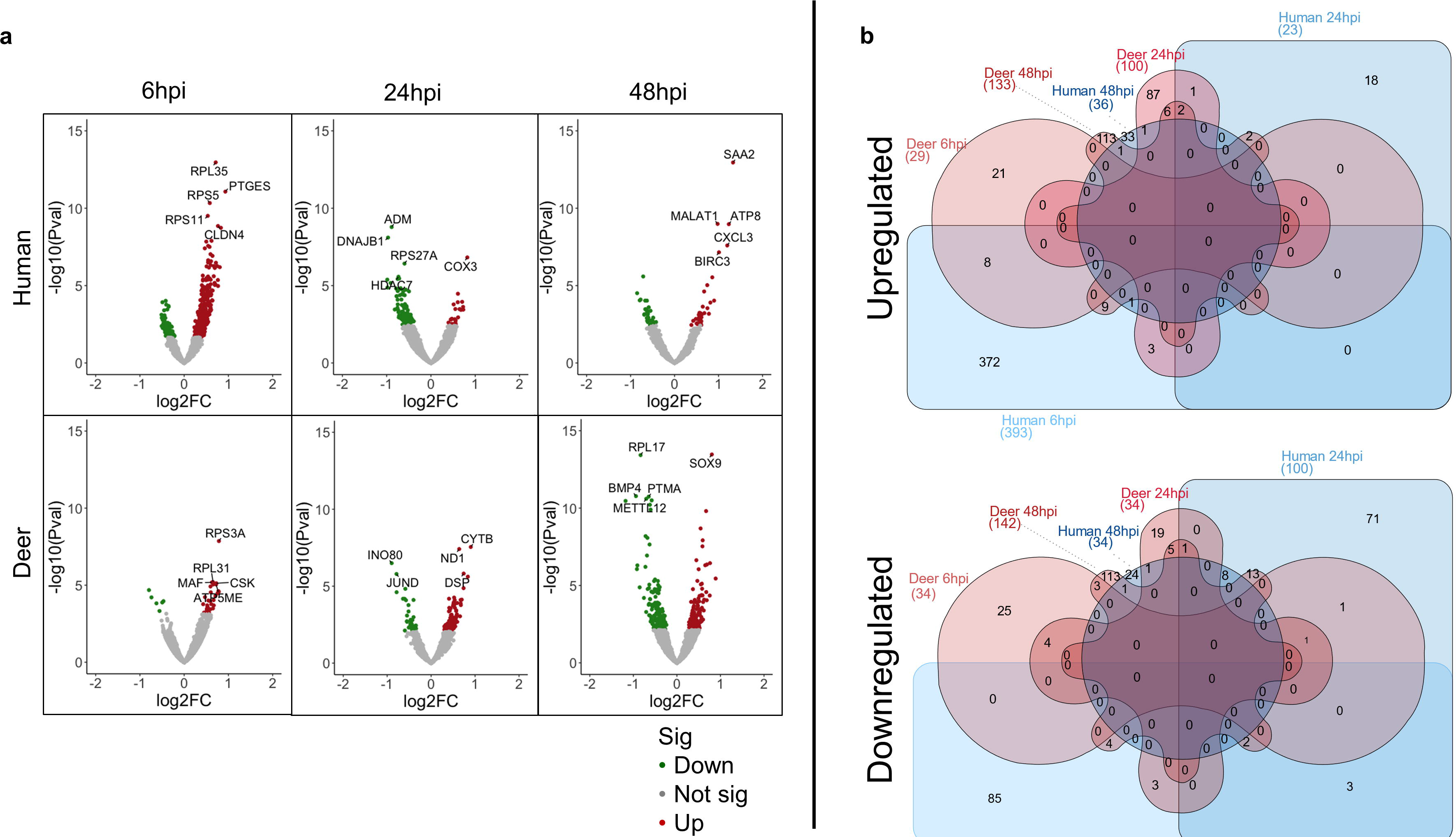
Graphical representation of differential gene expression in Deer-RECs and HRECs infected with SARS-CoV-2. (a) Volcano plots showing differential gene expression (DEG) at each time-point (6, 24, 48 hpi) in Deer-RECs (Deer) and HRECs (Human). Upregulated genes were shown in red, downregulated in green, and grey as differentially expressed but statistically insignificant. The top five gene genes were labeled as determined by the *P* value and fold change at each time point. b) Multidimensional six-set Venn diagrams showing significantly expressed DEGs shared between time points (6, 24, 48 hpi) in Deer-RECs (Deer) and HRECs (Human). Upregulated and downregulated genes were plotted separately to avoid any confounding genes that may be significantly expressed at three-time points or in both species but with a differential directional fold change between those time points or species. Shapes in red represent deer samples, while blue represents human samples. Data represent three technical replicates.

### Comparative pathway enrichment in HRECs and Deer-RECs infected with SARS-CoV-2

There were more significant (*P* < 0.05) enriched pathways in HRECs compared to Deer- RECs (Supplementary Data). At 6 hpi, 470 pathways were significantly enriched in HRECs compared to the 96 in Deer-RECs, and 93 of these pathways were significant (*P* < 0.05) in both. Among these shared pathways, the IL-17 signaling pathway was one of the most significant pathways observed in the Deer-RECs. At 24 hpi, 340 pathways were significant (*P* < 0.05) in HRECS, while 303 were significant in Deer-RECs and 243 of these pathways were significant (*P* < 0.05) in both. In addition to the IL-17 signaling pathway, the pathogen-induced cytokine storm signaling pathway was also significant (*P* < 0.05) in both HRECs and Deer-Recs. At 48 hpi, 410 pathways were significant (*P* < 0.05) in HRECS, while 222 were significant (*P* < 0.05) in both species, including the IL- 17 signaling and Pathogen Induced Cytokine Storm signaling pathways.

#### IL-17 signaling pathway

SARS-CoV-2 inoculation of HRECs and Deer-RECs cultures triggered contrasting signaling events in the IL-17 pathway at 6 hpi (Fig. 3). Deer and human cells showed clear divergence in activating early signaling genes such as *HSP90* and predicted regulation of *TRAF3IP2, TRAF5, TRAF2,* and *SRSF1*, leading to the difference in mRNA stabilization. Although *HSP90* showed no differential expression in Deer-RECs, it was significantly downregulated (*P* < 0.05) in HRECs (-0.41 log_2_FC), with enhanced expression of *MAP2K2* (0.52 log_2_FC)*, RELA* (0.42 log_2_FC) and predicted activation of *NF*κ*B- p50*, and *CEBP*β. A predicted activation of *NF-*κ*B* in SARS-CoV-2 infected HRECs was shown to influence the predicted upregulation of several genes associated with proinflammatory cytokine response, chemoattraction (*CCL2, CCL11, CCL20, CXCL2, CXCL5, CXCL8/IL8*), and hypersecretion of mucus (*MUC5AC, MUC5B*).

**Figure 3.**
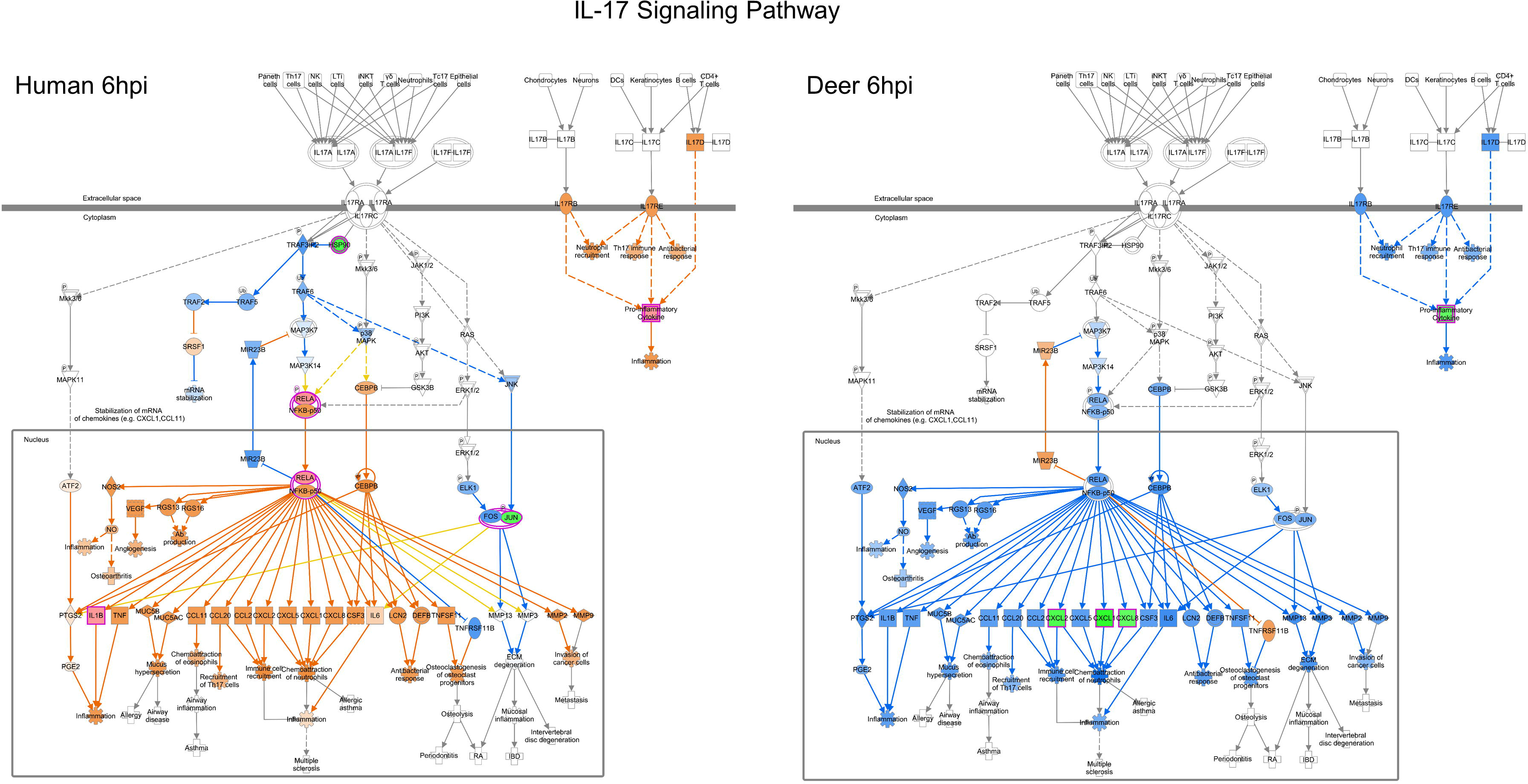
Differential gene expression within the canonical IPA IL-17 signaling pathway for both human and deer respiratory epithelial cells at 6hpi. The Ingenuity Pathway Analysis software (Qiagen) molecule activity predictor tool was used to predict the IL-17 signaling pathway activity based on significant differential gene expression. All significant DEGs were outlined in magenta, and a double outline indicates the involvement of multiple genes. Upregulated genes were shown with pink fill, while downregulated genes were shown with green fill. Similarly, predicted activation is shown in orange, and predicted inhibition is shown in blue. The greater the up/down expression or more confident the prediction, the darker the fill. Predictions that were inconsistent with the state of the downstream molecule were shown in yellow. Solid lines represent direct relationships, while dashed lines represent indirect relationships. Data represent three technical replicates.

Indeed, there was an increase in *IL1*β expression (0.41 log_2_FC) at 6 hpi in SARS-CoV-2 infected HRECs (Supplementary Data). In contrast, SARS-CoV-2 infected Deer-RECs showed a robustly predicted downmodulation of the proinflammatory cytokine and chemokine response, as evidenced by < -0.7 log_2_FC in the expression of *CXCL1, CXCL3,* and *CXCL8* (Supplementary Data).

#### Cytokine Signaling pathway

A significant down-modulation of the cytokine *TNF* and chemokines *CXCL3* and *CXCL8* was observed in SARS-CoV-2 infected Deer-RECs within the first 24 hpi. Interestingly, the NF-κB inhibitor *NFKBIA* or *I*κ*B*α (-0.51 log_2_FC), and *SOCS3* (-0.51 log2FC), a negative feedback regulator in cytokine signaling, were downregulated in Deer-RECs at 24 hpi (Fig. 4; Supplementary Data). In HRECs, *NGFR* (-0.75 log_2_FC; in the TNF receptor complex), *SLC20A1*(-0.50 log_2_FC; in the glucose transporter complex), and *JUN* (-0.59 log_2_FC; in the AP1 complex) were downregulated at 24 hpi (Fig. 4). A group of genes associated with the NF-κB signaling pathway, i.e., *SAA2, TNFAIP3, BIRC3*, and *IRF1* were all upregulated in SARS-CoV-2 infected HRECs. By 48 hpi, an upregulation of *SAA2* (1.33 log_2_FC) and *TNFAIP3* (0.86 log_2_FC), both biomarkers of severe COVID-19 disease, was observed. The apoptosis inhibitor *BIRC3* was upregulated (0.38 log_2_FC) by 6 hpi, and expression increased two-fold (1.01 log_2_FC) in HRECs by 48 hpi. In addition, the expression of *IRF1* (0.61 log2FC) was also upregulated in SARS-CoV-2 infected HRECs at 48 hpi. The upregulation of NF-κB pathway associated gene coincides with a surge in *CXCL3* (1.20 log_2_FC) expression in SARS-CoV-2 infected HRECs at 48 hpi (Fig. 5). Meanwhile, in Deer-RECs, there was no differential regulation of genes associated with the NF-κB signaling, but an increase in the expression of *IFNAR* (0.41 log_2_FC)*, CXCL6* (0.69 log_2_FC), and *CXCL8* (0.57 log_2_FC) was observed at 48 hpi.

**Figure 4.**
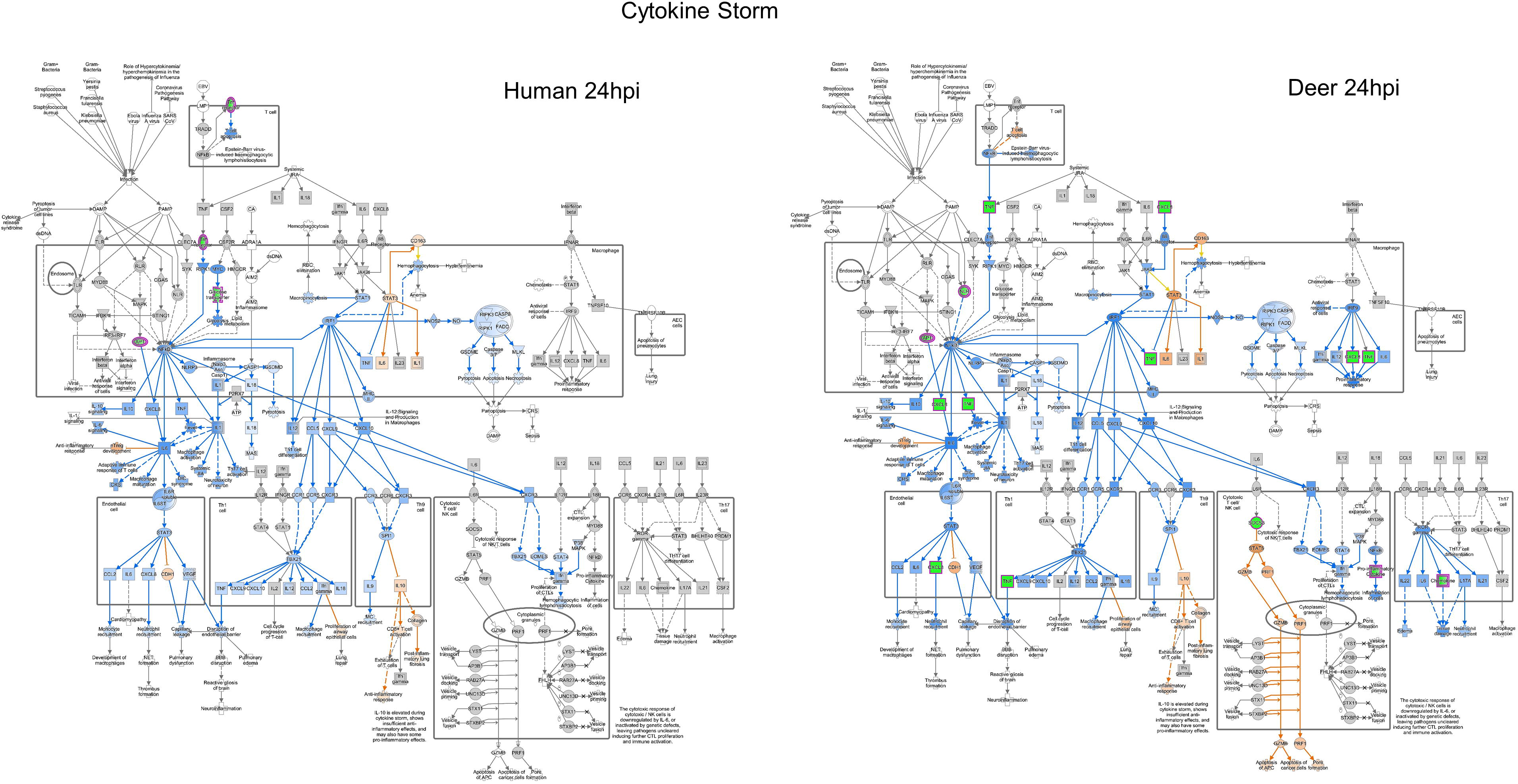
**Differential gene expression within the canonical IPA pathogen-induced cytokine storm signaling pathway for both human and deer respiratory epithelial cells at 24hpi**. The Ingenuity Pathway Analysis software (Qiagen) molecule activity predictor tool was used to predict the pathogen-induced cytokine storm signaling pathway activity based on significant differential gene expression. All significant DEGs were outlined in magenta, and a double outline indicates the involvement of multiple genes. Upregulated genes were shown with pink fill, downregulated genes were shown with green fill, and genes with grey fill had fold changes close to zero. Similarly, predicted activation is shown in orange, and predicted inhibition is shown in blue. The greater the up/down expression or more confident the prediction, the darker the fill. Predictions that were inconsistent with the state of the downstream molecule were shown in yellow. Solid lines represent direct relationships, while dashed lines represent indirect relationships. Data represent three technical replicates.

**Figure 5.**
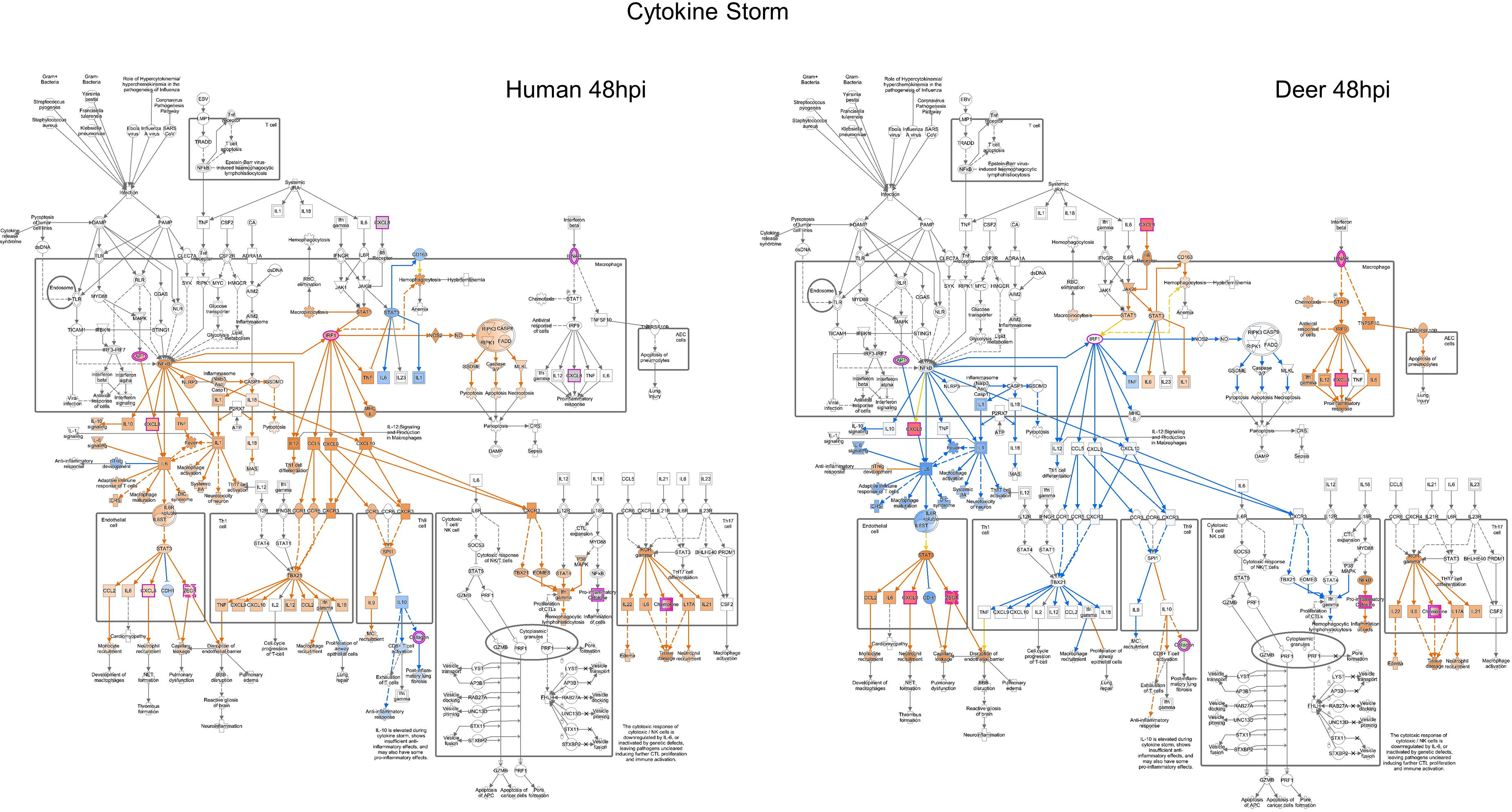
Differential gene expression within the canonical IPA pathogen-induced cytokine storm signaling pathway for both human and deer respiratory epithelial cells at 48hpi. The Ingenuity Pathway Analysis software (Qiagen) molecule activity predictor tool was used to predict the pathogen-induced cytokine storm signaling pathway activity based on significant differential gene expression. All significant DEGs were outlined in magenta, and a double outline indicates the involvement of multiple genes. Upregulated genes were shown with pink fill, downregulated genes were shown with green fill, and genes with grey fill had fold changes close to zero. Similarly, predicted activation is shown in orange, and predicted inhibition is shown in blue. The greater the up/down expression or more confident the prediction, the darker the fill. Predictions that were inconsistent with the state of the downstream molecule were shown in yellow. Solid lines represent direct relationships, while dashed lines represent indirect relationships. Data represent three technical replicates.

## DISCUSSION

An intriguing question about COVID-19 disease is how SARS-CoV-2 interplays with the host during infection, such that the virus causes a broad clinical spectrum of disease in humans whereas it replicates and transmits readily, yet infection remains subclinical in WTD. Studies have shown that deer lung cells (ATCC CRL6195) ^8^ and human bronchial/tracheal cells (ATCC PCS-300-010) are susceptible to SARS-CoV-2 ^21^. This is the first study to infect primary respiratory epithelial cell cultures derived from these two mammalian species and perform a comparative transcriptomic analysis over the course of the infection with SARS-CoV-2 to identify virus and host cell molecular factors responsible for the different clinical outcomes reported in human and deer. The findings presented here could aid in identifying common, perturbed gene networks that outline a shared/divergent host targetome for SARS-CoV-2 and provides biomarker candidates for targeted drug design.

The current study established that Deer-RECs and HRECs were susceptible to SARS-CoV-2 infection, and the virus replicated in these cells in an infectious dose-dependent manner as evidenced via observation of virus-induced CPE, detection of viral nucleoprotein in infected cells by immunostaining and viral sequence alignments obtained from the transcriptomic analysis. Particularly, SARS-CoV-2 mediated cell death was greater in Deer-RECs compared to HRECs. Our previous study showed that SARS-CoV-2 induces cytotoxicity rather than apoptosis in HRECs ^21^. The present RNA-seq analysis in HRECs further supports a more than two-fold increase in the expression of *BIRC3* from 6 to 48 hpi, suggesting that SARS-CoV-2 inhibits apoptosis at a very early stage of infection (Supplementary Data). Based on HRECs and Deer-RECs morphological changes, i.e., cell death (nuclear fragmentation) and DEG data, there was no indication of Deer-RECs undergoing apoptosis. Furthermore, the downmodulation of *HMGB1* (-0.48 log_2_FC) (Supplementary Data) by 48 hpi in Deer- RECs, a marker associated with necrotic cell death ^27^, also suggested that cell death may not be necrotic. Therefore, the mechanism of cell death in SARS-CoV-2 infected Deer-RECs should be further investigated.

The RNA quality obtained after the virus and mock-inoculation at each time point in both cell types was high, with almost no degradation ^28^. The obtained sequence reads were aligned to the *Homo sapiens* GRCh38.p13 (HRECs) and *Odocoileus virginianus* 1.0 (Deer-RECs) as reference assemblies, respectively. An important limitation to address was that the human reference genome has been much more robustly annotated than the WTD reference genome. However, WTD transcripts aligned with humans with an average identity of 89.79% and average coverage of 86.60% ^29^. This limitation was explicit during IPA analysis, as there were a significantly higher number of pathways detected in humans (470, 340, and 410) compared to WTD (96, 303, and 222) at 6, 24, and 48 hpi, respectively. Nevertheless, the analysis and discussion of all the pathways differentially regulated in Deer-RECS and HRECs was beyond the scope of this study. Rather, the discussion of the present study focuses mainly on the cytokine signaling pathway since the most severe clinical presentation resulting from SARS-CoV-2 infection in humans has been associated with a sudden acute increase in circulating levels of different proinflammatory cytokines (“cytokine storm”) ^16, 17^.

### Early innate immune mediators associated with SARS-CoV-2 entry in HRECs and Deer-RECs

SARS-CoV-2 upregulated *ATP6AP1* (0.32 log_2_FC) gene expression at 6 hpi in HRECs (Supplementary Data). ATP6AP1 is involved in membrane trafficking, and SARS-CoV-2 non-structural protein 6 directly interacts with this protein, leading to impaired lysosomal acidification in lung epithelial cells ^30, 31^. SARS-CoV-2 uses a non-lytic lysosomal egress pathway for virus release ^32^. In contrast, no significant differential expression of *ATP6AP1* was observed at 6 hpi in Deer-RECs; however, the expression of the serine incorporator (SERINC) transmembrane protein family *SERINC2* was significantly upregulated at 6 hpi. In 2022, Timilsina and others ^33^ reported that SERINC5 and SERINC3 restricted SARS-CoV-2 entry in lung epithelial cell lines. Additional studies are required to establish the role of *SERINC2* and SARS-CoV-2 entry in Deer-RECs.

### Early innate immune mediators associated with IL-17 could be responsible for divergence in cytokine signaling in human and deer respiratory epithelial cells

Cumulative evidence from both *in vitro* studies using human airway epithelial cells and clinical cases have shown the role of early signaling pathways, including the cytokine storm signaling, coronavirus pathogenesis, influenza A signaling, and IL-17 signaling pathway ^34–37^. Indeed, the IL-17 signaling pathway was among the most significantly enriched pathways identified in the present study.

Interestingly, genes such as *GRO1*/*CXCL1*, *CXCL3,* and *CXCL8* that regulate the IL-17 and cytokine storm signaling pathways in human SARS-CoV-2 infections ^34^, were upregulated in HRECs but downregulated in Deer-RECs as early 6 hpi. SARS-CoV-2 ORF3a, M, ORF7a, and N viral proteins activate NF-κB and induce proinflammatory cytokine expression ^38^. Epigenetic and single-cell transcriptomic analyses have demonstrated that NF-κB transcription is essential for SARS-CoV-2 replication ^39^. A possible factor influencing the divergence in the modulation of the NF-κB expression between the human and deer cells observed at 6 hpi could be the predicted differential expression of miRNA *MIR23B*, which was downregulated in the HRECs and upregulated in Deer-RECs at 6 hpi (Fig. 3). In fact, MIR23B plays a key role in IL-17, NF-κB, MAPK, and IFN-associated pathways and RIG-mediated signaling pathways ^40, 41^. The miR23b is predicted to bind SARS-CoV-2 ORF1ab and has high expression in human lungs ^42^, and it is known to hinder human rhinovirus HRV-1B infection by decreasing the very low-density lipoprotein receptor ^43^. A study by Pierce and others ^42^ reported that miRNA was a key differentiating factor between SARS-CoV-2-resistant and -susceptible cells. That is, miR23b was among the most differentially upregulated miRNA enriched in the resistant cells and after 24 hpi its expression was significantly downregulated in susceptible cells ^44^. Alterations of the miRNA expression in epithelial cells may contribute to the pathogenesis of chronic and acute airway infections. Hence, analyzing the role of these types of small noncoding RNA in antiviral immune responses and the characterization of miRNA target genes might contribute to a better understanding of the mechanisms of interplay between the host and viruses toward developing therapeutic strategies for the prevention and treatment of acute SARS-CoV- 2 infection.

The AP-1/JUN is a single transcription factor that regulates various cellular processes, including cell proliferation, differentiation, and apoptosis ^45^. The ability of a single transcription factor to control a collection of biological processes makes it an attractive target for signal transduction modification by viral proteins. The N protein of SARS-CoV was found to activate the AP1 pathway as a strategy to regulate cellular signaling ^46^.

Most recently, the spike protein of SARS-CoV-2 has been reported to induce JUN transcription via MAPK activation ^47^, leading to increased IL-6 release, which has been proposed as a mechanism for the initiation of hyper-inflammatory response, cytokine storm, and multi-organ damage associated with severe cases of SARS-CoV-2 infection ^47^. In the present study, there was a sustained suppression of *JUN* genes in HRECs up to 24 hpi, and an increase in expression activation was predicted by 48 hpi. Contrarily, in the Deer-RECs, *JUN* expression was significantly downregulated at 48 hpi, resulting in the predicted inhibition of IL-6. The induction of IL-6 is a key contributor to cytokine storm signaling, and a simultaneous increase along with STAT3 can amplify signaling machinery for an exacerbated inflammatory response also involving the NF-κB signaling pathway ^18^. Even though no significant *IL-6* upregulation was observed in SARS-CoV-2 infected HRECs, there was an increase in the expression of *STAT3* at 6 hpi (Supplementary Data), leading to a predicted activation of *IL-6* (Fig. 3). SARS-CoV-2 target the upstream mediators of the Jak-STAT pathway to impair interferon signaling across several human cell types ^48^

Another transcription factor of Jak-STAT signaling associated with IL-6 production during virus infection is SOCS3 ^49^, a negative feedback regulator in cytokine signaling, which also plays an important role during apoptosis, inflammation, T-cell development, and viral infection ^50^. The presence of SOCS3 reduces the induction of various types of IFNs, and in turn, a delayed IFN response can result in early viral spread leading to pulmonary and systemic inflammation in critical cases of SARS-CoV-2 ^51^. Moreover, using SOCS1/3 antagonists can block the replication and release of SARS-CoV-2 in human lung cell lines ^51^. While there were no changes in the expression of *SOCS3* in HRECs over the course of infection, Deer-RECs showed significant downregulation in response to SARS-CoV-2 infection by 24 hpi. A hypothesis that should be considered is whether a decrease in *SOCS3* expression in Deer-RECs at 24 hpi followed by an increased expression of *IFNAR1* (Fig. 5) at 48 hpi could be an innate immune response to abrogate viral replication and release. The significant downregulation of *LAMP1* observed at 48 hpi in Deer-RECs further supports this, which is in line with a previous study reporting that an increase in the expression of the *LAMP1* gene promoted SARS- CoV-2 production via exocytosis in Vero cells ^52^.

### Transcriptional activation of the NF-**κ**B signaling pathway is a critical innate immune response between human and deer respiratory epithelial cells

The SARS-CoV-2 N protein triggers the hyper-activation of NF-κB by undergoing liquid- liquid phase separation, which recruits kinases TAK1 and IKK. Furthermore, the inhibition of the liquid-liquid phase separation of the N protein has been shown to restrict NF-κB activation essential in virus-induced dysfunctional inflammatory responses and cytokine storm ^53^.

In this study, the increased expression of transcription factors *RELA* and *STAT3* at 6 hpi in HRECs led to the predicted activation of the expression of NF-κB associated cytokines *IL6* and *TNF* and chemokines *CCL2, CCL11, CCL20, CXCL1, CXCL2, CXCL5, and CXCL8/IL8*. The upregulated expression of *IL1*β further supports the differential modulation observed at 6 hpi (Fig. 3; Supplementary Data). The simultaneous upregulation of *RELA/p65* and *STAT3* by 6 hpi in HRECs could be a possible factor for amplifying cytokine signaling, as previously reported by Hojyo and others ^18^. Dysregulated NF-κB signaling pathway in HRECs continued into 48 hpi, with evident upregulation of NF-κB signaling factors such as *SAA2, TNFAIP3, BIRC3*, and *IRF1* ^54, 55^. The activation of these molecules is predicted to disrupt the epithelial barrier, inhibit the proliferation of airway epithelial cells, and activate the inflammasome ^38^.

In contrast, in SARS-CoV-2 infected Deer-RECs, the transcription factors *RELA* and *NF-*κ*B* signaling were predicted to be inhibited by 6 hpi, and this prediction continued through 48 hpi. Consequently, the expression of *CXCL1, CXCL3,* and *CXCL8* were suppressed along with predicted downregulation of other chemokines (*CCL2, CCL11, CCL20, CXCL5*) and cytokines (*IL1*β*, IL6, TNF, CSF3*). Although NF-κB activity predicted in Deer-RECs was similar to HRECs at 24 hpi, it is clear *SOCS3* or *NFKBIA*/*I*κ*B*α, an NF-κB inhibitor ^56^, might play a role in abrogating the expression of *TNF* and *CXCL8* in Deer-RECs. Similarly, TNF has also been shown to induce inflammatory cell death and lead to lethal cytokine shock in mice ^15^. The ability of Deer- RECs to downregulate *CXCL8* and *TNF* expression early may be key to their ability to weather the cytokine storm associated with SARS-CoV-2 infection. Single-cell transcriptomic analysis of bronchial lavage fluid from SARS-CoV-2 infected human patients with severe symptoms, including a surge in inflammation and airway damage, revealed higher expression of CXCL8, resulting in neutrophil infiltration of the lungs, causing lung epithelial damage ^57^. *CXCL8* has also been identified as a hub gene in the process of SARS-CoV-2 through protein-protein interaction network analysis, emphasizing its key role in SARS-CoV-2 infection ^58^. Among many roles, CXCL8 is also involved in tissue repair by promoting the migration and proliferation of cells ^59^.

*MAP4K4,* which is involved in regulating cell migration and invasion ^60^, was only upregulated at 48 hpi, along with *CDKN1A/p21*, which has a role in cell proliferation by regulating DNA replication and repair ^61^. At the same time, in Deer-RECs, there is also an upregulation of *VEGFA* (0.60 log_2_FC) and *HBEGF* (0.34 log_2_FC) (Supplementary Data), which are associated with enhancing compensatory lung growth through paracrine signaling ^62^. However, in human cases, the upregulation of serum biomarkers VEGFA and HBEGF was associated with the severity of SARS-CoV-2 infection ^63^.

Nonetheless, CXCL8, MAP4K4, p21, VEGFA, and HBEGF are necessary for wound repair, and their collective and delayed upregulation at 48 hpi in Deer-RECs may be associated with tissue repair signaling.

## CONCLUSION

The comparative transcriptomic data analysis in human and deer primary respiratory epithelial cells infected with SARS-CoV-2 cells reported herein support that the dysregulation of IL-17 and NF-κB signaling pathway could be one of the major drivers for the divergent early innate immune response between these mammalian species. The findings from this study could be extrapolated to explain the lack of clinical signs reported in WTD under experimental and field conditions as opposed to severe clinical outcomes in humans affected by SARS-CoV-2. Additional research is necessary regarding the deer “omics” and SARS-CoV-2, however, due to the scarcity of BSL3 facilities for large animals, utilizing 3D cell cultures of WTD as an alternative approach can potentially improve our comprehension of host-virus interactions, ultimately resulting in innovative intervention approaches.

## MATERIALS AND METHODS

### Isolation of white-tailed deer respiratory epithelial cells (Deer-RECs)

The state wildlife veterinarian provided aseptic trachea sections from hunter-killed WTD. In brief, samples from the mid to lower tracheal region (6-8 inches) were aseptically collected in Dulbecco’s Minimum Essential / Ham’s F-12 medium with GlutaMAX (DMEM/F-12) (Thermo Fisher Scientific, Waltham, MA, USA), supplemented with 100 IU/mL of penicillin/100 µg/mL of streptomycin (Pen-Strep; Thermo Fisher Scientific), and 1.25 µg/mL of amphotericin B (AmpB) (Thermo Fisher Scientific) for isolation of Deer- RECs as previously described ^21^, and transported to the laboratory soon after the field dressing. Samples were washed and incubated in phosphate-buffered saline (PBS) pH 7.4 supplemented with Pen-Strep to remove blood clots. Then, samples were incubated at 4°C for 48 h in a digestion medium [calcium and magnesium-free Minimum Essential Medium (MEM; in-house), supplemented with 1.4 mg/mL pronase (Millipore-Sigma, Burlington, MA, USA), 0.1 mg/mL DNase (Millipore-Sigma), 100 µg/mL Primocin^®^ (Invivogen, San Diego, CA, USA)]. Tissue digestion was neutralized using 10% heat-inactivated fetal bovine serum (EqualFetal FBS; Atlas Biologicals, Fort Collins, CO, USA). The tissue digest containing cells was passed through a 100 µm cell strainer, washed, pelleted, and resuspended in DMEM/F12 medium. Collected cells were either seeded directly using respective growth medium or frozen in LHC^®^ basal medium (Thermo Fisher Scientific) containing 30% FBS and 10% dimethyl sulfoxide (DMSO) (Millipore-Sigma).

### Culture of primary deer (Deer-RECs) and human respiratory epithelial cells (HRECs)

Isolated primary Deer-RECs and commercially acquired HRECs (ATCC, Manassas, Virginia, USA; PCS-300-010, Lot-70002486) were subcultured on cell/tissue culture flasks or plates (Greiner Bio-One North America Inc, Monroe, NC, USA), pre-coated with PureCol® Type I collagen (40 µg/mL/4 mm^2^; Advanced BioMatrix, Inc., San Diego, CA, USA). Both Deer-RECs and HRECs were propagated in growth media [ATCC airway epithelial cell basal medium (ATCC^®^ PCS-300-030™) supplemented with bronchial epithelial cell growth kit (ATCC^®^ PCS-300-040™), Pen-Strep and Amp-B (Thermo Fisher Scientific)]. Cells were subcultured by dissociation with 0.5 X TrypLE^TM^ express enzyme (Thermo Fisher Scientific) and neutralized using 50% heat-inactivated FBS, mixed in LHC basal medium. Primary cells used in this study corresponded to passage 3 for Deer-RECs and passage 9 for HRECs.

### SARS-CoV-2 propagation and titration *in vitro*

SARS-CoV-2 (BEI Resources, SARS-Related Coronavirus 2, Isolate USA-WA1/2020, NR-52281) was propagated in Vero-E6 cells (ATCC, CRL-1586) according to previous protocols ^21, 22^. In brief, cells were sub-cultured in DMEM (Corning, Tewksbury, MA, USA) supplemented with 10% FBS incubated at 37°C in humidified 5% CO_2_ incubator. Cell debris-free viral supernatants were collected from SARS-CoV-2 virus inoculated culture flasks showing cytopathic effect (CPE) in ≥90% of Vero-E6 cells. Virus titration by plaque assay ^23^ resulted in a stock titer of 10^7^ PFU/mL. Virus stock was aliquoted and frozen at -80°C for subsequent virus infectious studies on HRECs and Deer-RECs.

### SARS-CoV-2 infections in HRECs, and Deer-RECs

For virus titration assays, ∼20,000 cells (Vero-E6/ HRECs/ Deer-RECs) per well were seeded in a 96-well plate (CellBIND Costar; Corning) and, 24 h prior to infection, the cells were washed once with LHC medium and pre-incubated with an infection medium containing ATCC airway epithelial cell basal medium, 2% Ultroser-G (Sartorius Stedim Biotech GmbH, Goettingen, Germany), 1X 4-(2-hydroxyethyl)-1- piperazineethanesulfonic acid (HEPES) (Thermo Fisher Scientific), 1X MEM non-essential amino acids (Thermo Fisher Scientific), 1X Glutamax (Thermo Fisher Scientific), Pen-Strep and AmpB (Thermo Fisher Scientific). For transcriptomic analysis, 6-well plates with a seeding density of 300,000 cells per well on the day of infection were washed once with LHC medium and inoculated with infection medium containing different doses of SARS-CoV-2 (10^5^, 10^4^, 10^3^, 10^2^, 10, 1 PFU/mL) or mock inoculated with infection medium only. After 2 h incubation at 37°C and 5% CO_2_, the inoculum was removed, cells were washed once with LHC medium, replaced with fresh infection media, and incubated for 6, 24, and 48 h. Following infection, virus-induced CPE, including rounding of cells, cell detachment, clumping, and dead cells, were recorded. For imaging, the cells on plates were fixed in 4% paraformaldehyde (Electron Microscopy Sciences, Hatfield, PA, USA).

### Immunocytochemistry staining in Deer-RECs and HRECs

Immunocytochemistry staining (ICC) was used to confirm the expression of the SARS-CoV-2 nucleocapsid (N) protein as described previously ^21^. In brief, 4% paraformaldehyde-fixed cells were permeabilized with 0.1% Triton X-100 (Millipore- Sigma) for 10 min. Cells were blocked with animal-free buffer (Vector Laboratories, Newark, CA, USA) for 30 min and incubated overnight (16 h) at 4°C with a recombinant rabbit anti-SARS-CoV-2 N protein monoclonal antibody (0.75 μg/mL) [BEI Resources, Monoclonal Anti-SARS Coronavirus/SARS-Related Coronavirus 2 Nucleocapsid Protein (produced *in vitro*), NR-53791; SinoBio Cat: 40143-R001]. The cells were treated with 0.1% hydrogen peroxide for 5 min, followed by 1 h incubation with ImmPRESS VR anti-rabbit IgG HRP polymer detection kit (MP-6401-15; Vector Laboratories). Chromogenic detection *in situ* was performed using ImmPACT DAB EqV peroxidase substrate solution (Vector Laboratories) and counterstained with hematoxylin. Microscopic images were captured using an Olympus® CKX4 microscope (Olympus^®^ Corp., Center Valley, PA, USA), Infinity 2 camera, and Infinity Analyze imaging software (Ver 6.5.5, Lumenera Corp, Ottawa, ON, Canada).

### RNA Extraction and Sequencing

For transcriptome analysis, cells were lysed using TRizol reagent (Thermo Fisher Scientific), and total RNA was isolated from cells after performing the chloroform phase separation, followed by purification with MagMax Total RNA Kit (Thermo Fisher Scientific). According to manufacturer instructions, RNA quality was assessed with a 2100 Bioanalyzer system (Agilent Technologies, Santa Clara, CA, USA). Library preparation was performed with the 3’ QuantSeq kit, and 100 bp single-end reads were generated utilizing the Illumina Hiseq 6000. Library preparation and sequencing were performed at the Iowa State University DNA Facility (Ames, IA, USA).

### Differential Gene Expression Analysis

Tools present at galaxy.scinet.usda.gov were utilized to analyze the sequenced reads. FastQC and MultiQC were used to perform quality control for reads and examine raw read data and counts. Trim Galore! (version 0.6.7) was used to remove adapters and reads with a phred score below 20. HiSat2 (version 2.1.0) ^24, 25^ was used to align the trimmed sequence to the *Homo sapiens* GRCh38.p13 and *Odocoileus virginianus* 1.0 assemblies, respectively. Raw counts were generated with FeatureCounts (version 2.0.1). Differential gene expression (DEG) was performed using DeSeq2 (version 2.11.40.6) ^26^ utilizing a parametric fit type and poscounts to account for genes with zero counts. DEG analysis was based on the model treatment + hpi + treatment:hpi +E for each species. Significant DEGs were reported for the interaction effect of treatment:hpi for each species and were declared statistically significant at a Benjamini-Hochberg False Discovery Rate (FDR) of 0.15. Gene names are based on Ensembl identifications. The *Odocoileus virginianus* reference genome is currently poorly annotated, and 75 significant DEGs (Supplementary Data) from the Deer-RECs did not have an Ensembl gene name. The Fasta sequence for each of these transcripts was input into the Blastn suite. Annotated genes with 95% or greater sequence identity were identified for 63 of these transcripts, and 6 were identified as long noncoding RNAs (lncRNAs) (Supplementary Data). The 63 annotated genes were utilized in subsequent analysis.

### SARS-CoV-2 genome alignments

Sequences that did not align to the *Homo sapiens* GRCh38.p13 or *Odocoileus virginianus* 1.0 assemblies, respectively, were written out into separate files and were subsequently aligned to the SARS-CoV-2 reference genome ASM985889v3 using Bowtie2. The number of SARS-CoV-2 genome alignments for each sample was graphed using GraphPad Prism 9.5.0 (GraphPad Software Inc., La Jolla, CA, USA).

### Pathway Analysis

The DEG data were analyzed using Qiagen Ingenuity Pathway Analysis (IPA) software (Qiagen Digital Insights, Redwood City, CA, USA) to identify the significantly enriched IPA canonical pathways differentially regulated in HRECs and Deer-RECs over the course of the infection. Specifically, the canonical pathway function of IPA core analysis was used to identify significantly enriched canonical pathways from the lists of DEGs following inoculation of HRECs and Deer-RECs with SARS-CoV-2 at 6, 24, and 48 hpi.

Based on the right-tailed Fisher’s Exact Test, canonical pathways were declared significant at *P* < 0.05. The molecule activity predictor (MAP) tool was used to predict the upstream and downstream effects of activation and inhibition based on these known changes in gene expression. These pathways were utilized to view DEG divergences between species at each time point.

## DATA AVAILABILITY

The data underlying this article are available in the article and its online supplementary material. Please contact the corresponding authors for any additional data.

## SUPPLEMENTARY DATA

Supplementary Data are available online.

## ACKNOWLEDGEMENTS

The authors thank Dr. Rodger Main for his generous support in obtaining various SARS- CoV-2 reagents from Biodefense and Emerging Infections (BEI) Research Resources Repository. We appreciate the contributions and technical expertise of Sarah Anderson. The following reagents NR-52281, and NR-53791, were deposited by the Centers for Disease Control and Prevention and obtained through BEI Resources, NIAID, and NIH. Lastly, we wish to acknowledge the licensed hunters that contributed fresh deer tissues used in this study

## AUTHOR CONTRIBUTIONS STATEMENT

Resources, supervision and funding acquisition, and project administration by LGL, LCM. RKN and LGL led the project, procured reagents from BEI resources, designed experiments, curation and analysis of generated data and visualization, and drafted the original manuscript. Deer samples were provided by RR. RKN performed cell cultures and immunostaining, including validation. KSP performed all BSL3 experiments, such as SARS-CoV-2 infections. Isolation of RNA, RNA quality, Transcriptomic analysis, software, and pathway analysis was performed by KMSD. BB is in charge of all the BSL3-related work. BB co-led the project and assisted in experimental design. Data curation analysis and presentation RKN and KMSD. Manuscript original draft preparation RKN, KMSD, LCM, YS, LGL. Manuscript review and editing KMSD, RKN, LCM, LGL, YS, KP, BB, RR.

## SUPPLEMENTARY INFORMATION

HREC and Deer-REC DEGs, Deer gene annotations, enriched IPA pathways were listed in supplementary data spreadsheet.

